# Symmetry breaking in the embryonic skin triggers a directional and sequential front of competence during plumage patterning

**DOI:** 10.1101/491092

**Authors:** Richard Bailleul, Carole Desmarquet-Trin Dinh, Magdalena Hidalgo, Camille Curantz, Jonathan Touboul, Marie Manceau

**Author notes:** To whom correspondence should be addressed; these authors contributed equally to this work.

## Abstract

The development of an organism involves the formation of patterns from initially homogeneous surfaces in a reproducible manner. Simulations of various theoretical models recapitulate final states of natural patterns^1-4^ yet drawing testable hypotheses from those often remains difficult^4,5^. Consequently, little is known on pattern-forming events. Here, we extend modeling to reproduce not only the final plumage pattern of birds, but also the observed natural variation in its dynamics of emergence in five species. We built a unified model intrinsically generating the directionality, sequence, and duration of patterning, and used *in vivo* experiments to test its parameter-based predictions. We showed that while patterning duration is controlled by overall cell proliferation, its directional and sequential progression result from a pre-pattern: an initial break in surface symmetry launches a traveling front of increased cell density that defines domains with self-organizing capacity. These results show that universal mechanisms combining pre-patterning and self-organization govern the timely emergence of the plumage pattern in birds.

## INTRODUCTION

The diverse shapes and motifs that adorn animals have been a long-standing interest of developmental biologists and theoreticians: how can patterns arise from homogeneous structures during the development of an organism in an often highly organized and reproducible manner? Numerous modeling studies, frequently assuming a chemical basis for pattern-forming factors (for review^1,2^) but also recently integrating cellular and mechano-chemical processes (for review^3,4^), led to the theorization of self-organizing dynamics to explain the emergence of many patterns, guiding in a few cases the identification of candidate factors *in vivo*. However, a given final pattern can often be reproduced by a variety of models^4,5^, and frequently this reproduction is only approximate. In addition, each of these models (stationary solutions and equations) putatively corresponds to many developmental mechanisms. Therefore, in the absence of spatial reference allowing for candidate approaches, choosing and building models to predict and test *in vivo* patterning mechanisms has remained challenging.

The plumage pattern is one of the few emblematic systems in studies of pattern formation and evolution in which the modeling-experimentation gap has been successfully bridged^5,6^. In birds, feathers are implanted in so-called “tracts” (or *pterylae*) separated by glabrous areas that are thought to allow steric space necessary for flight movements^7^ (Figure 1). The spatial distribution of tracts at the scale of the whole body (i.e. macro-pattern) is broadly conserved, all birds having capital (head), humeral / alar (wings), dorsal, ventral, femoral / crural (legs) and caudal (tail) tracts^8^. However, their shape and size, as well as the geometrical arrangement of feathers within tracts (i.e., micro-pattern) vary between bird groups (as formerly studied in the zoological field of pterylography;^9,10^).

**Figure 1:**
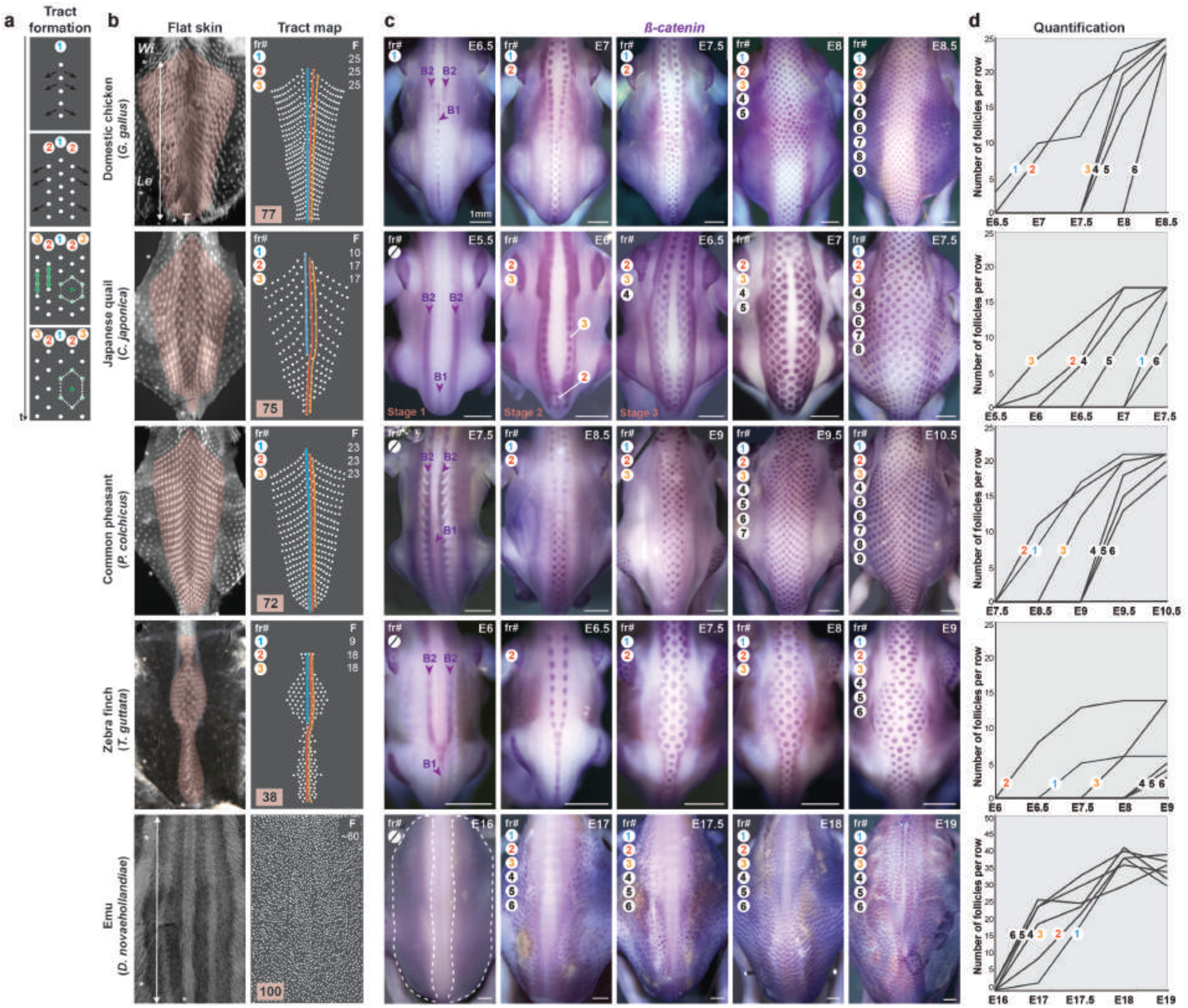
Dorsal tract formation varies in directionality, sequentiality and duration. **(a)** During development, the dorsal tract appears in a medial-to-lateral bi-directional wave (black arrows) of feather rows (fr#1-3 are schematically represented), resulting in a micro-pattern in which each feather follicle (dark green) is surrounded by a regular hexagon of neighbors (light green). This pattern later distorts along the antero-posterior axis (green arrows). **(b)** Flat preparations of dorsal skins (left panels) and their corresponding schematic map (right panels) show that completed dorsal tracts (in pink) vary in size between the domestic chicken (where it covers ∼77% of the dorsum surface; n=2), the Japanese quail *C. japonica* (75%; n=3), the Common pheasant *P. colchicus* (72%; n=3), and the zebra finch *T. guttata* (38%; n=2). In these four species, feather follicles (in white) organize in longitudinal rows (fr; circled numbers) that extend from the neck to the tail and contain a reproducible number of feathers F (counted from wings to tails as shown by white arrows): F=25, 10, 23, and 9 for fr#1 (in blue; note that in the Japanese quail and the zebra finch, this row is shorter as it is composed of late developing feathers^10,17^) and F=25, 17, 23, and 18, for fr#2 and fr#3 (in orange and red). In the emu *D. novaehollandiae* (n=3) the tract covers 100% of the dorsum surface and at E26 we quantified ∼F=60 feathers along tract length). **(c)** *In situ* hybridizations revealing *β-catenin* transcripts (in purple) mark the dynamic appearance of feather rows at equivalent developmental stages. In the domestic chicken, the Japanese quail, the common pheasant, and the zebra finch, the first feather rows arise from three medial bands: one posteriorly located (B1) and two anteriorly located (B2). Lateral rows then appear one by one in a front that progresses ventrally and stops at the limit of the dorsal tract. In the emu, *β-catenin* is first seen in two large surfaces flanking a thick medial band (white dotted line); follicles then individualize progressively at random locations. **(d)** Automatic quantifications of F per row show that fr#1-6 appear in a sequential manner in the first four species. Follicle individualization occurs in a given row when the preceding row is near completed (with the exception of fr#1 where it has species-specific timing and fr#2 in the quail where it is first restricted posteriorly). In contrast, along virtual longitudinal lines in the emu (see Supp. Figure 1) follicles individualize with no apparent spatial order. They invade the feather field in 2-3 days in Galliforms and the zebra finch, and 1 day in the emu. **E**, embryonic day; **t**, time; stars show the position of wings (**Wi)**, legs (**Le**) and tails (**T**); Stage 1, Stage 2 and Stage 3 are defined in Figure 3 and 4; scale bars: 1mm.

Work performed in the domestic chicken *Gallus gallus* showed that the patterning of feather tracts involves surface dynamics occurring in the developing skin tissue. In this species, the early mesodermal layer first directs the formation of competent epidermal areas (or feather fields). Within each feather field, feather follicles individualize in a medial-to-lateral “wave” of differentiation to form a regular dotted pattern, which later becomes distorted to give rise to a reproducible geometry in which each feather is surrounded by an elongated hexagonal array of neighbors^7^ (see Figure 1a). Feather arrangement is thus a complex yet ordered motif that results from the timely orchestration of patterning events in the developing skin including (1) local individualization of shapes (here, follicles) from an initially homogeneous feather field, and (2) the directional and gradual progression of this differentiation process. Efforts to tackle patterning factors have largely relied on theoretical approaches. Computer simulations of self-organizing models (e.g., reaction-diffusion, chemotaxis), alone or in combination, can give rise to regularly spaced dots reminiscent of the chicken feather motif^11,12^ (or other cutaneous structures in Vertebrates^13,14^) which suggested plumage pattern results from a self-organization of the developing skin. This work helped identifying candidate cellular events (epidermal/dermal interactions^12^) and molecules (including Shh, BMPs, and FGFs^11,12,15,16^) involved in the individualization of feather follicles. However, models did not explain the directional and progressive aspects of tract formation.

To tackle this challenge, we extended the predictive power of modeling to reproduce not only the tract pattern, but also the common and varying attributes of both final tracts and of their dynamics of emergence we identified in five species of birds. We built a model (combining parameters of reaction-diffusion, chemotaxis, and local cell proliferation) that independently generates the observed spatio-temporal attributes of patterning without addition of an extrinsic mathematical wave. We varied parameters of this unified model, and tested *in silico* results using *in* and *ex vivo* experimentation. We found that *(i)* spatial heterogeneities present before the appearance of feather follicles launch the bi-directional patterning wave, *(ii)* the latter travels as a front of increased cell density that defines domains with self-organizing capacity and controls pattern sequentiality, and *(iii)* overall cell proliferation rate controls the duration of tract completion.

## RESULTS AND DISCUSSION

We characterized the spatial organization of the plumage in the dorsal region of five bird species. The chicken *Gallus gallus* historically served as model: consistent with previous work^7^ we observed that when completed (at E11) its dorsal tract covers ∼77% of the skin surface and displays an orderly arrangement in which adjacent longitudinal rows (i.e., feather rows; fr) contain a reproducible number of feather follicles (F=25 as quantified between wings and tail in medial rows) and form “chevrons” along the dorso-ventral axis. For comparison, we chose two close relatives in the Galliform bird group, namely the Japanese quail *Coturnix japonica* and the common pheasant *Phasianus colchicus* in which we previously described feather distribution^17^. In Galliforms, overall tract size (∼75 and 72% of dorsum surface, respectively), shape, and geometry are conserved, while the number of feathers per row is species-specific: F=17 in the quail (except in fr#1 known to develop later where F=10^17^) and F=23 in the pheasant. In the zebra finch *Taeniopygia guttata*, a passerine bird in which tracts have been previously thoroughly described^10^, the dorsal tract encompasses a relatively thinner skin region (∼38% of the dorsum surface) compared to Galliforms. It displays a different shape with an enlargement at the level of hind limbs (so-called “saddle”), with fewer and shorter rows that are less strictly arranged (9<F<18 in the medial rows). Finally in the emu *Dromaius novaehollandiae*, which is part of the derived Ratite group of flightless birds, we observed feathers across the whole dorsum in a dense and disorganized pattern (F∼60 as quantified along lines drawn from wings to tail; Figure 1a,b). These observations show that independently of dorsum size, both the macro-pattern (relative tract surface and shape) and the micro-pattern (number, spacing and geometry of feather follicles) vary between species.

To link tract patterns to events involved in their formation we compared the dynamics of feather follicle emergence. We used stains for *β-catenin* transcripts that mark the early differentiation of feather follicles^18^ and developed an automatic algorithm to consistently sort and count the number of follicles in microscopy images (Supp. Figure 1). We found that in all Galliforms and in the zebra finch, *β-catenin* initially forms one visible medial (B1) and two lateral (B2) longitudinal bands, in which follicle individualization gradually takes place, starting at the posterior end of the band and progressing anteriorly. Adjacent feathers rows form according to the same steps and sequentially, in a row-by-row wave that travels ventrally and stops at the limit of the feather field, with most follicles in a given row having formed prior to the appearance of the next *β-catenin* expressing band. Follicles reach the limit of the tract in ∼2/3 days, irrespective of the duration of the whole development (varying from 14 days in the zebra finch to 16-22 days in Galliforms). Thus, tract patterning is characterized by medial-to-lateral bi-directionality, sequence (i.e., row-by-row dynamics and individualization prior to next row) and duration broadly conserved between the four bird species. These spatio-temporal attributes may thus controled by shared developmental mechanisms. We however observed subtle variation in the initial *β-catenin* pattern: while B2 bands are similar in all species, the medial band B1 extends from the inter-limb region to the tail in the chicken while in the Japanese quail it is reduced to the posterior region, in the common pheasant it is diffuse and in the zebra finch it is fused to lateral bands B2, forming a Y-shape. This profile defined the species-specific location of first-individualizing follicles, suggesting a link between the initial organization of the feather field and the timely dynamics of follicle differentiation.

In contrast with the first four species, *β-catenin* covers the whole feather field except a thick medial band in the emu. Follicle individualization occurs randomly in space and time, covering the surface defined by initial *β-catenin* expression pattern in ∼1 day (emus develop in 60 days; Figure 1c,d). This absence of directionality and sequence for follicle appearance in a comparably shorter duration may be due to entirely different patterning mechanisms specific to Ratites, or to changes in the processes that control the dynamics of tract establishment independently of those governing follicle individualization.

To guide the identification of patterning factors, we built a model recapitulating both shared and varying characteristics of tract patterning in the different species. Reaction-diffusion dynamics, first theorized by Alan Turing^19^ and involving the diffusion of at least one local self-activating factor *u* and its longer-range inhibitor *v*^19,20^ have been widely used to explain the establishment of natural patterns^1,2^. We simulated a range of reaction-diffusion equations for various interaction functions *f* and *g* on homogeneous surfaces as well as on an initial longitudinal line (as *β-catenin* initially forms medial bands in Galliforms and the zebra finch):

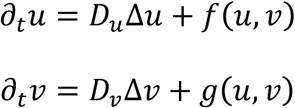

Consistent with previous studies, we found that such models can generate dotted patterns, but fail to reproduce the row-by-row sequence observed experimentally (Supp. Figure 2). This argues for the involvement of another mechanism orchestrating the sequential emergence of follicles. Key candidate mechanisms are cell proliferation (local cell density *n* evolving according to a proliferation rate *p*) combined with chemotaxis, the process of cell migration in response to a chemo-attractant *u* generated by the cells (at a rate *α*_*u*_), and also subject to diffusion (diffusivity *D*_*u*_) and degradation (rate *δ*_*u*_):

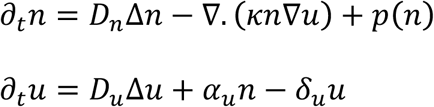

**Figure 2:**
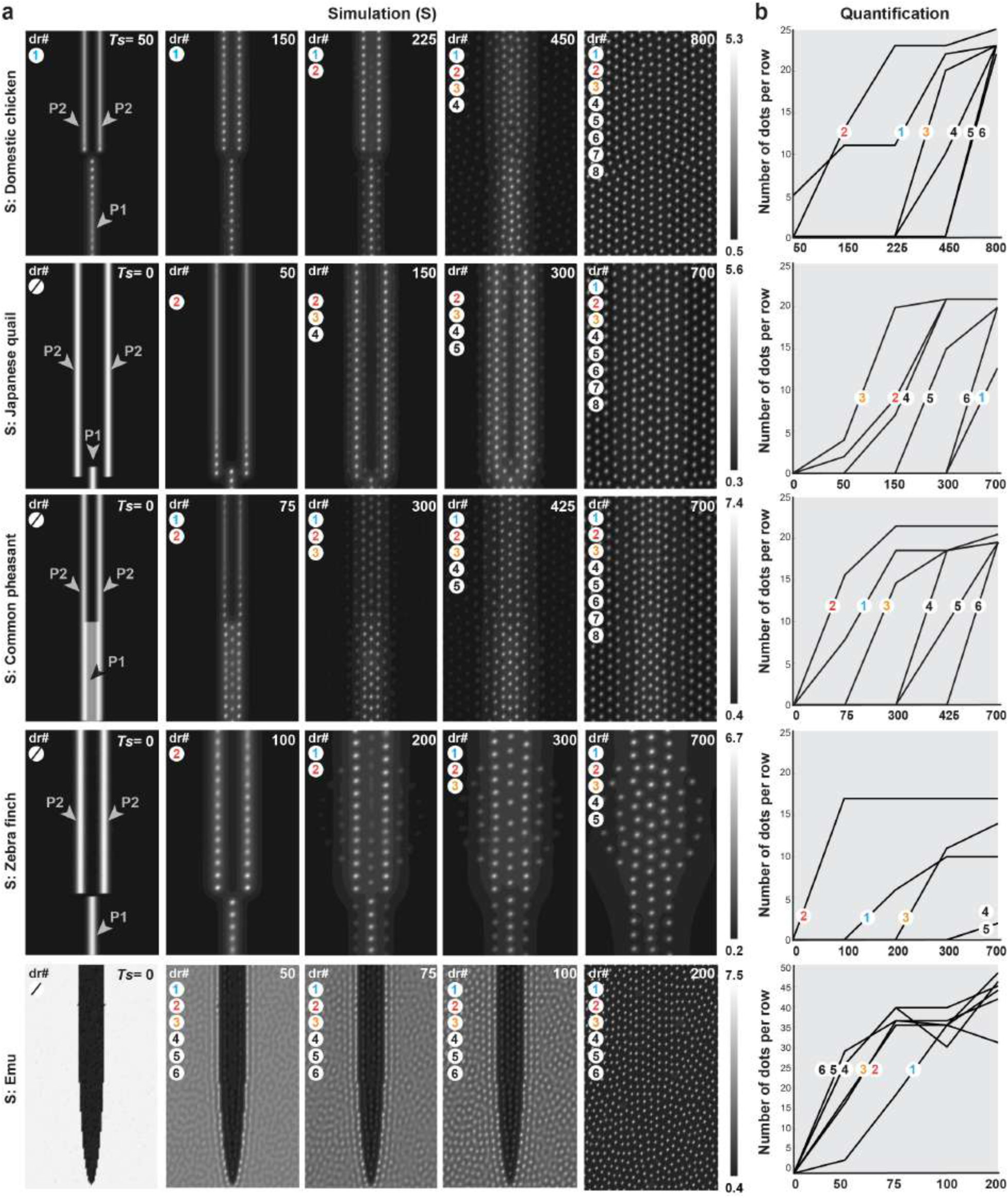
Simulations of the unified model reproduce spatio-temporal attributes. **(a)** *In silico* simulations (S) of the unified model were run on frames corresponding to (1) the relative size of the dorsal tract in each species and (2) numerical initial conditions that define one posterior (P1) and two anterior (P2) longitudinal bands (left panels) respectively corresponding to *in vivo* measurements of B1 and B2 in the chicken, the Japanese quail, the common pheasant, and the zebra finch, and to two surfaces separated by a thick medial band in the emu (see Figure 1). At various intermediate simulation times (*Ts*; indicated in white), dots individualize in rows. The formation of dotted rows (dr) occurs in a bi-directional and sequential manner in the first four sets of simulations, and randomly in the fifth (mimicking *in vivo* species-specific dynamics). Simulations stabilize with motifs corresponding to the final macro and micro-patterns observed in respective species. For each species-specific simulation color bars indicate minimal-to-maximal *n* values in a black-to-white gradient. **(b)** Automatic quantifications of dots at various simulation times confirm that the unified model reproduces both shared directionality/sequentiality in tract formation and inter-species variation in follicle number.

Such models create dotted motif^21^ but do not allow the formation of spots in a sequential manner, and when applied to an initial longitudinal line, we found that they produce transient but not stable longitudinal bands; Supp. Figure 3). Thus, alone, and even when forced onto axial initial conditions, self-organization models do not recapitulate the highly orchestrated dynamics establishing tracts observed *in vivo*.

**Figure 3:**
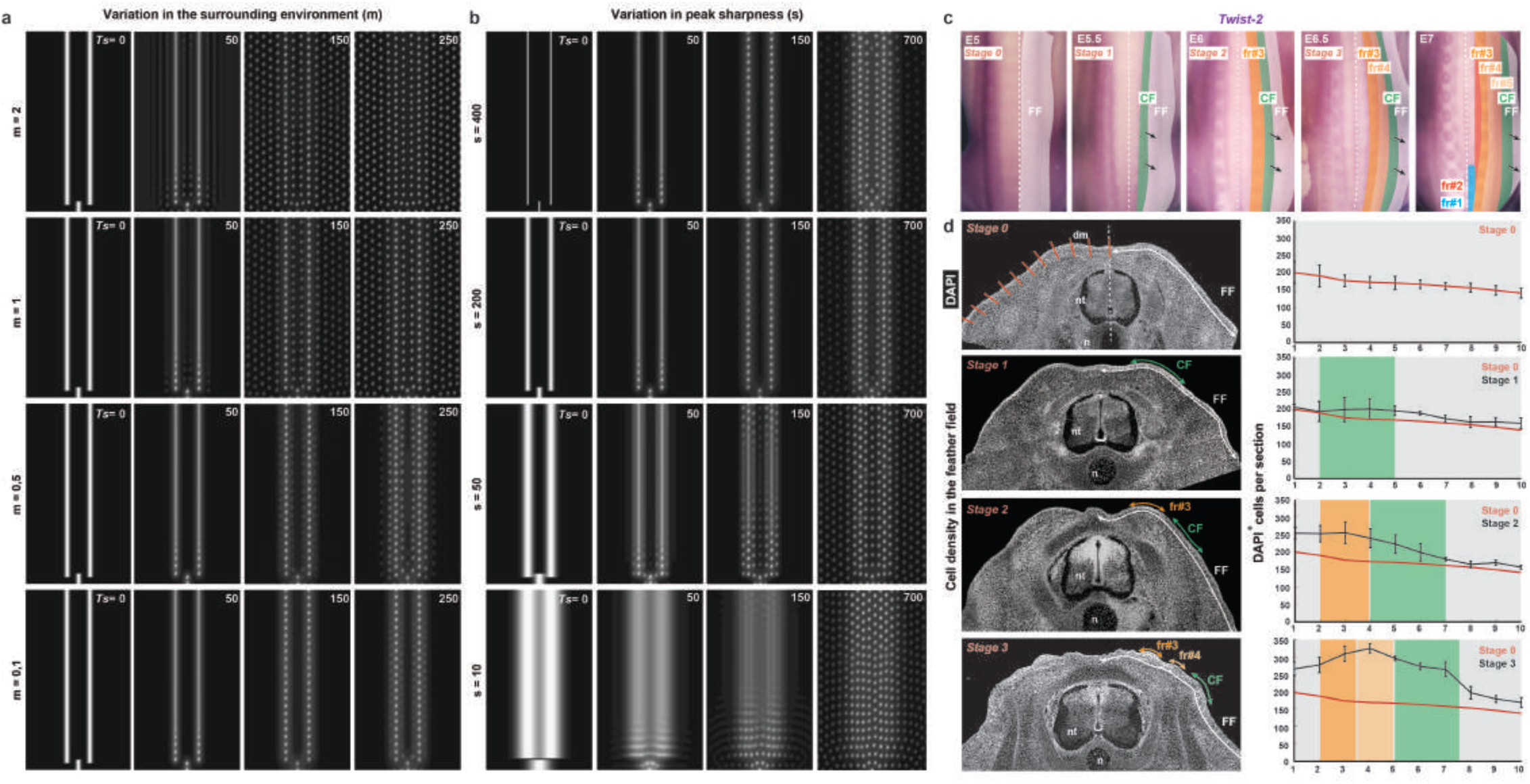
Sequentiality results from the progressive appearance of longitudinal domains. **(a)** Simulations of the unified model with initial conditions corresponding to the Japanese quail show that dots form in a row-by-row sequence when the minimal density outside of Gaussian peaks *m* is low, but simultaneously when *m*>0,5. **(b)** Increasing peak width **s** leads to a loss of row-by-row sequence while decreasing *s* has no effect on pattern emergence. **(c)** *In situ* hybridizations revealing *Twist-2* transcripts (in purple) from E5 (Stage 0) to E7 mark the progressive appearance of longitudinal domains progressing (black arrows) in a competence front (CF; in green) within the feather field (FF; in white) and later forming feather rows (i.e., at that level in the quail, fr#3 and fr#4 that form first, shown in shades of orange, followed by fr#2, fr#1, fr#5 respectively in red, blue, and light orange). **(d)** Left panels: local changes in cell density can be observed on transverse sections of Japanese quail embryos stained with DAPI to mark nuclei (in white) prior to *β-catenin* expression (Stage 0), at initial conditions of *β-catenin* expression (Stage 1), and during the individualization of the fr#3 and fr#4 (Stages 2 and 3). Right panels: quantifications of skin cells within 10 sections corresponding to the red bars shown at Stage 0 show that cell density gradually increases in a traveling medial-to-lateral wave (results obtained at Stage 0 are shown in dark red in all other graphs for comparison). **Ts**: Simulation time; **E**, embryonic day; **nt**: neural tube; **n**, notochord.

Temporal sequences were previously generated when chemotaxis and molecular interactions are driven through external forcing: dots form in a chicken-like sequence with a model that responds to a medial-to-lateral wave of cell “competence”^11^. Recently, a model of epidermal/dermal interaction also forced onto a medial-to-lateral priming wave produced condensed structures in a sequence of directionality and speed related to the properties of the wave^12^. Similar modeling in other study systems (e.g., sensory organs in Drosophila) also yielded sequential patterning^22^. To uncover factors providing directionality and sequentiality to follicle individualization, we thus developed a partial differential equation model containing self-organizing mechanisms, but devoid of extrinsic spatio-temporal forcing. This model describes local cell density *n*(*t, x*) evolution according to diffusion, chemotaxis towards high concentrations of an activator chemo-attractant, and intrinsic logistic proliferation accounting for a division rate *α*_*n*_ at low cell population levels and a carrying capacity of the tissue *β*_*n*_, density above which proliferation stops. The instantaneous concentration of the activator *u*(*x, t*) evolves according to its autocatalytic production by cells and the concentration of its repressor *v* (*x, t*).

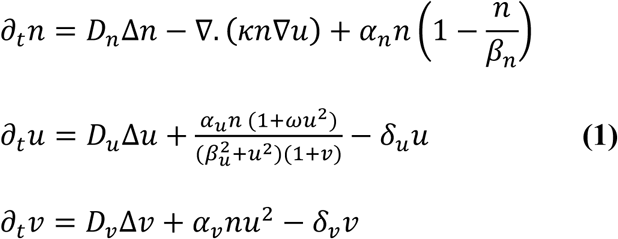

*D*_*n*_, *D*_*u*_ and *D*_*v*_ parameterize the diffusion of cells, attractor and repressor, respectively, while *κ* accounts for the sensitivity to chemotaxis, *α*_*u*_ (*α*_*v*_) is the production rate of the attractor (repressor) by the cells, *β*_*u*_ and *ω* respectively quantify the saturation threshold and autocatalysis sensitivity of the activator, and *δ*_*u*_ (*δ*_*v*_) is the degradation rate.

As expected from the properties of both reaction-diffusion and chemotaxis systems alone, this unified model allows the formation, from an initially homogeneous expression profile, of regularly spaced dots that appear throughout the simulation surface in a timely fashion but at random locations (Supp. Figure 4), consistent with tract formation observed in areas of the emu skin marked by *β-catenin*. To reproduce the observed variation in pattern emergence we thus adapted the size of numeric simulation frames to that of tract size/shape in each bird and built geometrical initial conditions for simulations corresponding to the initial expression pattern of *β-catenin* in all species: we used three Gaussian equations restricted along the axes corresponding to the measured central location of B1 and B2 *β-catenin* bands relative to body landmarks (i.e. *x*_2_ and *y*_1_, *y*_2_, respectively; Supp. Figure 5):

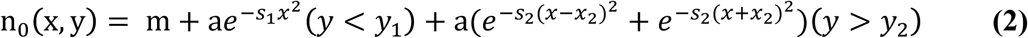

In these equations, *m* represents a minimal density throughout the simulation frame and *a, s*_1_ and *s*_2_ are the amplitude and sharpness of medial (P1) and lateral (P2) peaks, respectively (Supp. Figure 6). We found that simulations of the model with a unique set of parameters but species-specific initial conditions allow the formation of individualized dots in a bi-directional and typical row-by-row sequence requiring similar times of simulation *Ts* to stabilize in a final pattern (*Ts* = 700/800) for Galliforms and the zebra finch, and in a spatially random and faster manner (*Ts* =200) in the emu (Figure 2). Together, the incremental building of a unified model reproduced shared and varying spatio-temporal attributes of tract pattern emergence in all species, allowing us to predict that these attributes rely on an early spatial organization of the yet un-patterned skin as marked by the expression of *β-catenin*.

To characterize this initial organization, we generated model-based predictions by varying parameters defining Gaussian peaks (corresponding to longitudinal bands *in vivo*) and their surrounding environment (corresponding to the feather field). We used simulations with Japanese quail-like initial conditions, which are the simplest. We found that varying *a*, which represents here the amount of causal factor in longitudinal bands, does not affect the final pattern nor its dynamics of formation (Supp. Figure 7). Thus, patterning occurs independently of the initial amount of cells and/or molecules responsible for follicle differentiation within bands. It is however impacted by the amount of factors outside of peaks, described here by *m*: proper row-by-row sequence is lost when *m* reaches a high threshold value. Strikingly however, we found that sequential patterning occurs with low *m* values (close to *m*=0; Figure 3a). Thus, conserving low minimal density of cellular/molecular factors outside of spatially restricted domains along the medial axis is an initial break in symmetry that endows only the regions within these domains with the capacity to form patterns (neighboring regions having close to basal density unable to generate patterns). Even when minimal, this break in symmetry acts as a trigger, sufficient to launch the medial-to-lateral patterning wave in which cells near longitudinal peaks progressively transmit the pattern-forming capacity to neighboring regions.

To understand what controls patterning sequentiality, we tested whether the row-by-row dynamics results from self-organizing events (as they have been previously involved in intrinsic properties of patterning waves^11,12^). To do so, we varied all self-organization parameters. We observed changes in the size and spacing of dots when, all parameters otherwise equal, we modified the activator or repressor diffusion *D*_*u*_ or *D*_*v*_ (Supp. Figure 8), the chemotaxis sensitivity κ (Supp. Figure 9), the strength of molecular interactions *α*_*u*_, *α*_*v*_, *β*_*u*_ and *ω* (Supp. Figure 10), or the degradation rate *δ*_*u*_, *δ*_*v*_ (Supp. Figure 11). Except for extreme values often causing an absence of individualization, all tested values largely preserved a row-by-row dynamics of dot emergence. These simulations suggest that self-organization controls follicle size and spacing consistent with previous findings^5,12,23^, but have no impact on the onset of the row-by-row wave nor are responsible for its sequential aspect. Interestingly, all simulations defined by species-specific frames and *β-catenin* profiles yielded a number of dots per row similar to that observed for follicles in each species (compare with Figure 1 and 2). The parameters of self-organizing events that control *in vivo* the individualization of follicles may thus be conserved, and we performed complementary simulations where these parameters are all maintained but the width of longitudinal bands containing causal factors is modified. We found that when we increased peak width (by decreasing *s*), several dot rows could form simultaneously, while a decrease (by increasing *s*) did not modify the resulting pattern or its sequential formation (Figure 3b). We therefore hypothesized that row-by-row sequentiality depends on the size of longitudinal domains possessing pattern-forming capacity. Consistent with this hypothesis, we found that in the Japanese quail the mesoderm and follicle marker *Twist-2* ^25^, though unlikely to contribute to the trigger wave as it is expressed throughout the feather field, delineates longitudinal surfaces appearing in the same order than future feather rows (Figure 3c). Competence within such spatially restricted domains is expected to impact local cell density, as condensation marks follicle formation^7,18,24^. We thus quantified cell density across the dorsal feather field of the Japanese quail, starting just before and at the “trigger stage” (i.e., corresponding to initial *β-catenin* expression; stage 0 and 1, respectively), and during the formation of the first formed rows (fr#3 and the posterior part of fr#2; stages 2 and 3, respectively). We found that at stage 0, cell density is slightly higher in the medial part of the dorsum and gradually decreases ventrally, not correlating with observed and simulated initial conditions. At stage 1 however, a peak of cell density appears at the level of initial *β-catenin* expression (i.e., of the putative first row). Its amplitude gradually increases, marking the differentiation of a follicle, as a second area of high cell density appears laterally (stage 2). The latter also increases and becomes flanked by a third peak at stage 3 (Figure 3d). Thus, a traveling wave of increased cell density progresses laterally. Together, simulations and experimental data indicate that row-by-row sequentiality results from a traveling front of longitudinal surfaces defined by causal factors in a sharp enough pattern that self-organizing parameters, controlling follicle size and spacing within these domains, authorize the formation of only one feather follicle at a time.

Because the timely appearance of surfaces with self-organizing capacity involves increased cell density, we next tested the effect of cell proliferation. We performed BrdU incorporation experiments on skin tissues at the trigger stage in the Japanese quail (in which tracts form sequentially and in 3 days). We found that BrdU^+^ cells (i.e., 20,3% of all DAPI^+^ cells) are homogeneously distributed in the feather field, showing proliferation does not convey symmetry breaking within the feather field (Figure 4a). We thus next varied the proliferation rate parameter *α*_*n*_ in a homogeneous manner throughout simulation frames. We found that varying *α*_*n*_ does not impact follicle size or spacing nor row-by-row sequence. However reaching final pattern states required longer simulation times as *α*_*n*_ decreased, and shorter simulation times as it increased, with a loss of row-by-row dynamics in the most extreme cases (Figure 4b). Together with BrdU incorporation experiments, these simulations suggested that cell proliferation controls the timing of tract completion. To test this prediction *in vivo*, we cultured explants of Japanese quail skin in varying concentrations of colchicine drug, known to inhibit cell proliferation. Low doses did not affect pattern compared to control experiments, while high doses had lethal effect (Supp. Table 1). We thus performed pulses of colchicine-mediated inhibition at the highest non-lethal dose and found that the speed of row formation reduced with longer colchicine pulses (identically to predictions observed with decreased *α*_*n*_). As a result, six days after treatment, tracts were not complete compared to control experiments (Figure 4c). Thus the proliferation rate of skin cells mediates the duration of the patterning process. This temporal attribute is comparable between Galliforms and the zebra finch irrespective of the duration of the whole development and despite variation in the relative size of their tracts. Proliferation rate may thus act as a constraint to the extent of feather fields that are characterized by different initial conditions, consistent with the observation that low values of *m* can also slightly modulate the timing of patterning (Figure 3).

**Figure 4:**
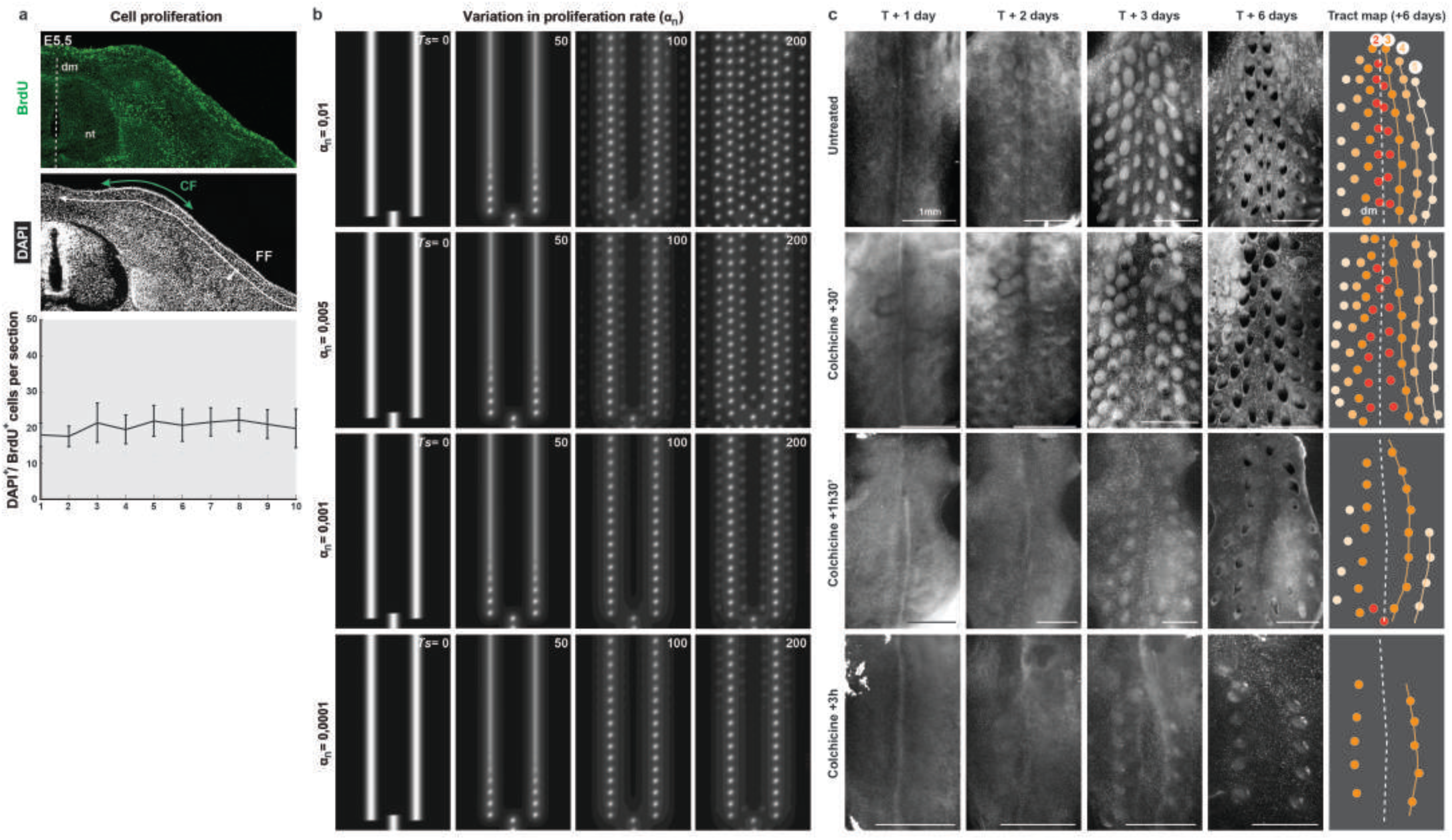
Cell proliferation controls the duration of tract formation. **(a)** Cell proliferation is revealed by BrdU stains (in green) on transverse sections of Japanese quail embryos at E5.5 (Stage 1/2) also stained with DAPI to reveal cell nuclei. Quantifications show that the proliferation rate is homogeneous throughout the feather field (i.e., BrdU^+^/ DAPI^+^ represent 20,3% of all DAPI^+^ cells on average in all counted sections as shown in Fig. 3; Friedman test, non significant). **(b)** Simulations of the unified model with initial conditions corresponding to the Japanese quail show that dot rows form through longer simulation times *Ts* (in white) when the overall proliferation rate *α*_*n*_ decreases. **(c)** The dynamic appearance of tracts occurs in a progressively slower manner in cultured explants of Japanese quail skins (prepared prior to the differentiation of the feather field) observed from one to six days (T+1 to T+6) after exposure to increasing pulses of colchicine treatment (from 30 minutes to 3 hours), compared to un-treated explants (schematics represent feather rows in the resulting tract with the color code used previously in Figure 1 and 3).

Altogether, simulations and experimental data allow us to propose a scenario in which tract patterning is a combination of reaction-diffusion, chemotaxis, and cell proliferation events taking place in a pre-patterned field. In Galliforms and the zebra finch, the distribution/behavior of *in vivo* factors (cells or molecules) controlling follicle differentiation (such as observed for *β-catenin*) is initially spatially restricted in longitudinal bands along the medial axis. This initial heterogeneity creates a break of symmetry in the surface field that triggers the appearance of longitudinal domains with self-organizing capacity. The medial-to-lateral progression of this competence front depends both on initial conditions of the system (resulting in its directional and sequential attributes), and optimal cell proliferation rate (yielding the completion of the patterning process in a timely fashion). In the emu, shallow initial conditions or high basal levels of causal factors (consistent with the observation that *β-catenin* is expressed throughout the dorsum surface in this species) do not sufficiently mark symmetry breaking, and even regions away from the peaks have the ability to self-organize, which results in larger competent areas and a loss of differentiation directionality.

This work sets the stage for the identification of patterning molecules, providing testable hypotheses on their origin (i.e., axial landmarks) and profile (i.e., correlation with initial conditions). Good candidates may comprise Wnt proteins^26^, which diffuse from axial tissues neighboring the skin (neural tube, somites) and activate the expression of *β-catenin*. It also highlights the importance of sequential aspects to patterning, which may explain how self-organizing mechanisms can act in a reproducible manner over large and developing surfaces.

## ACKNOWLEDGEMENTS

We thank B. Perthame for help with project conception and analyses, K. Painter, T. Lecuit, and F. Bosveld for helpful discussion, E. Bouvet, M. Ladner, N. Haupaix, and P. Galipot for help with specimen collection, and N. Quenech’Du and the Collège de France imaging facility for help with image processing and analysis.

## AUTHOR CONTRIBUTIONS

**RB** performed tract and expression analyses in Galliforms and the Emu, constructed the mathematical model and tested biological parameters *in silico* and *in vivo*. **CDT** conducted cell proliferation assays on skin explants. **MH** performed tract and expression analyses in the zebra finch. **CC** built the emu tract map. **RB**, **JT**, and **MM** designed the model and the experiments, developed the analysis tools and wrote the manuscript.

## MATERIALS AND METHODS

### Embryo sampling and flat skin preparation

Fertilized eggs were collected from a breeding colony at the Collège de France for zebra finches (*Taeniopygia guttata*), and from local suppliers for the other species: Les Bruyères élevage for domestic chicken *Gallus gallus*, Cailles de Chanteloup for Japanese quails *Coturnix japonica*, Les boix de Vaux for common pheasants *Phasianus colchicus*, and l’Emeu d’Uriage and Autruche de Laurette for emus (*Dromaius novaehollandiae*). After egg incubation in Brinsea Ovaeasy 190 incubators, embryos were treated *in ovo* with 9mg/mL of 5-Bromo-2’-deoxyuridine solution (Sigma; BrdU incorporation experiments), and dissected. Flat skins were prepared as described previously^17^. Specimens were fixed in 4% formaldehyde, and imaged.

### Expression analyses

*In situ* hybridization experiments were performed in each species (n is provided in Supp. Table 1) as described previously^27^ using antisense riboprobes synthesized from vectors containing 881-bp and 501-bp fragments respectively of Japanese quail and zebra finch coding sequences for *β-catenin* and a 740-bp fragment of Japanese quail coding sequence for *Twist-2*. Digoxigenin–labeled riboprobes were revealed with an anti-digoxigenin-AP antibody (1:2000, Roche) and an NBT/BCIP (Promega) substrate. Sequences of *β-catenin* primers are: F: AGCTGACTTGATGGAGTTGGA and R: TCGTGATGGCCAAGAATTTC (quail) F: TAGTTCAGCTTTTAGGCTCAGATG and R: CCTCGACAATTTCTTCCATACG (finch). Sequences of *Twist-2*primers are: F: AAAGCTCCAGTTCTCCTGTTTC and R: ATGTTGCTTCTCGCTTCTCTG.

### Quantifications of tract size and feather follicles/dots number

#### Tract size

Feather-containing surfaces normalized by that of the whole dorsum were measured using Fiji software on pictures of flat skins at developmental stages corresponding to tract completion (E11 for the domestic chicken, n=2; E10 for the Japanese quail, E12.5 for common pheasant, n=3, and E26 for the emu), and at hatching for the zebra finch (the relative surface of the completed tract being conserved during development, follicles are best visualized at P0; n=2).

#### Follicle/dot number

To quantify feather follicles or dot number (F) respectively on pictures of flat skins or model simulations in a time efficient and consistent manner (i.e, across species and at different stages), we developed a custom Matlab program. The algorithm follows three steps: first, a Gaussian filter reduces optical noise through image smoothing (Matlab function *imfilter*; filters were obtained with the function *fspecial*). Second, morphological operations are applied (i.e., closing followed by opening of the image with a disk of radius 1 pixel using functions *imopen* and *imclose*; the disk was constructed using the *strel* function). Third, feather follicles/dots are detected as connected components of the image and their properties (centroid, bounding box, surface, etc…) stored. Because background levels on skin images can fluctuate, the algorithm contains a set of linearly spaced thresholds to *(i)* produce a binary image through a detection threshold of the pre-treated image, *(ii)* identify the connected components of the binary image (Matlab function *bwconncomp*), and *(iii)* repeat the second step if the maximal value in a given component is above the next threshold.

The resulting feather/dot locations are presented in a Matlab interface (Supp. Fig. 1) allowing to adjust thresholds, filter properties, and correct for undetected follicles or false positives (needed for less than 5% of the total follicle number). Locations are processed to identify rows by segmenting the set of feathers/dots according to their location along the x-axis (and corrected by hand in the cases of Japanese quail and zebra finch for which fr# is shifted laterally and in the emu for which counting was performed along six virtual lines). Program code and interface (Dotfinder) are available on github.

### Modeling

All simulations were performed using FreeFem++^28^ software (specifically designed to compute numerical solutions of partial differential equations) with no flux (Neumann) boundary conditions (i.e., cells or molecules are free to diffuse outside of the tract). Spatial and temporal discretization parameters were chosen as a compromise between accuracy and efficiency (*n*_*x*_ between 120 and 160, *dt* between 0.01 and 0.1).

#### Size of simulations frames

For comparison of pattern dynamics relative to tract size, the width (l_s_) and length (L_s_) of simulation frames has been set for each species according to developmental landmarks: we measured widths (i.e., l, in mm, distance between wings) and lengths (i.e., L, in mm, distance between tails and a medial point between wings; Supp. Table 2 and Supp. Figure 5). We then adjusted coordinates for the domestic chicken so that the numbers of feather rows (i.e., fr=8) and of feathers per row (F=25) coincide *in vivo* and *in silico*. This allowed defining a 0.45 coefficient between L and L_s_ and a 0.7 coefficient between l and l_s_, which were reported to all other species. For better Figure readability, simulations were then rescaled to the same dimensions on all plots.

#### Initial conditions

Initial conditions for the emu were defined by a constant function times an indicator (which models the absence of signal at the midline) with small random fluctuations. For the other four species, they were implemented with equation **(2)** using parameters shown in Supp. Table 3.

#### Reaction-diffusion models

We tested a large number of reaction-diffusion models as shown in Supp. Table 4. Simulations in Supp. Figure 2 were performed using a model recently shown to reproduce a dotted pattern of denticles in sharks^25^:

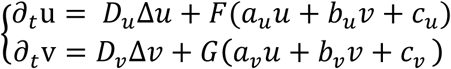

where *F* and *G* are rectifying functions avoiding negative or too large values of the argument: *F*(*x*) = *x* for 0 < *x* < *F*_*max*_, saturating at 0 and *F*_*max*_ outside of this interval, and *G*(*x*)= *x* for 0 < *x* < *G*_*max*_, saturating at 0 and *G*_*max*_ outside of this interval. Simulations were made on a square domain 0 < *x* < 75, 0 < *y* < 150, with parameters *D*_*u*_ = 0.02, *a*_*u*_ = 0.08, *b*_*u*_ = −0.08, *c*_*u*_ = 0.04, *d*_*u*_ = 0.03, *F*_*max*_ = 0.2, *D*_*v*_ = 0.6, *D*_*v*_ = 0.6, *a*_*v*_ = 0.16, *b*_*v*_ = 0, *c*_*v*_ = −0.05, *d*_*v*_ = 0.08, *G*_*max*_ = 0.5.

#### Chemotaxis models

We tested a large number of chemotaxis models such as described in^21^ Simulations in Supp. Figure 3 were performed using a model allowing the formation of a dotted pattern:

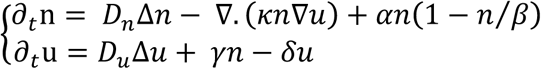

on a square domain 0 < *x* < 30, 0 < *y* < 60 with parameters *D*_*n*_ = 5, *D*_*u*_ = 0.1, *κ* = 5, *α* = 1, *β* = 3, *γ* = 1, *δ* = 1.

#### Unified model

All reference parameters are shown in Supp. Table 5 and were largely based on those used in a previous study of chicken-like plumage pattern formation^11^.

### Skin explants

Skin regions corresponding to putative dorsal tracts were dissected from E6 Japanese quail embryos and placed dermal side down on culture insert membranes (12-wells format, Falcon #353103) over 800 µL DMEM supplemented with 2% FCS and 2% Penicillin/Streptomycin. Stock solutions of Colchicine (50mg/mL in EtOH; Sigma #C9754) were diluted to various concentrations (0,00125 - 40 mg/mL) in the culture medium to identify the highest non-lethal dose (0,2mg/mL; Supp. Table 6). Pulse treatments were achieved by washing 0,2mg/mL Colchicine out in successive medium baths after 30 minutes (n=12), 90 minutes (n=7), or 3 hours (n=10), as opposed to untreated, control explants (n=9). Skin explants were incubated at 37°C with a 5% CO_2_ atmosphere (Thermo Scientific Midi 40); medium was changed every two days.

### Immuno-histological stains and quantifications

Embryonic specimens were embedded in gelatin/sucrose, sectioned using a CM 3050S cryostat (Leica), treated with HCl 2N for 20 minutes (for BrdU stains), rinsed and stained using a rat primary antibody directed against BrdU (Abcam; 1:200) and a Goat anti-rat Alexa 488 secondary antibody (Molecular Probes; 1:500). Cell nuclei were revealed using DAPI (Southern Biotech). Slides were mounted in Fluoromount (Southern Biotech) prior to imaging. For quantifications of cell density or proliferation, the number of DAPI^+^ or BrdU^+^ cells, respectively, was counted in ten sections of confocal images (defined from the dorsal midline to the ventral limit of the feather field using Fiji software; Figure 4). A student T Test calculated p values.

### Imaging

Flat skins and whole embryos were imaged using an AF-S Micro NIKKOR 60-mm f/2.8G ED macro-lens equipped with a D5300 camera (Nikon) and a MZ FLIII stereomicroscope (Leica) equipped with a DFC 450C camera (Leica). Confocal images were obtained using an inverted SP5 microscope (Leica) with a 40X immersed oil objective.

## REFERENCES

1. Kondo, S. & Miura, T. Reaction Diffusion as a framework for understanding biological pattern formation. Science 329, 1616–1620 (2010).

2. Green, J. B.A. & Sharpe, J. Positional information and reaction-diffusion: two big ideas in developmental biology combine. Development 142, 1203–1211 (2015).

3. Brinkmann, F., Mercker, M., Richter, T. & Marciniak-Czochra, A. Post-Turing tissue pattern formation: Advent of mechanochemistry. PLoS Comput Biol 14: e1006259 (2018).

4. Hiscock, T.W. & Megason, S. G. Mathematically guided approaches to distinguish models of periodic patterning. Development 142, 409 (2015).

5. Painter, K.J., Hunt, G., Wells, K., Johanneson, K. & Headon, D.J. Towards an integrated experimental–theoretical approach for assessing the mechanistic basis of hair and feather morphogenesis. Interface Focus 2, 433–450 (2012).

6. Baker, R. E., Schnell, S. & Maini, P. K. Waves and patterning in developmental biology: vertebrate segmentation and feather bud formation as case studies. Int. J. Dev. Biol. 53, 783–794 (2009).

7. Sengel, P. Morphogenesis of the Skin. Cambridge University Press(1975).

8. Neguer, J. & Manceau, M. Embryonic Patterning of the Vertebrate Skin. Reviews in Cell Biology and Molecular Medicine 3; 1 (2017).

9. Nitzsch, C.L. Nitzsch’s Pterylography. Journal of Anatomy and Physiology 2, 391 (1868).

10. Clench, M. H. Variability in Body Pterylosis, with Special Reference to the Genus Passer. The Auk 87, 50–691 (1970).

11. Michon, F., Forest, L., Collomb, E., Demongeot, J. & Dhouailly, D. BMP2 and BMP7 play antagonistic roles in feather induction. Development 135, 2797–2805 (2008).

12. Painter, K.J., Ho, W. & Headon, D.J. A chemotaxis model of feather primordia pattern formation during avian development. Journal of Theoretical Biology 437, 225–238 (2018).

13. Cooper, R.L. et al. An ancient Turing-like patterning mechanism regulates skin denticle development in sharks. Sci. Adv. 4, eaau5484 (2018).

14. Moustakas-Verho, J. E. et al. The origin and loss of periodic patterning in the turtle shell. Development 141, 3033–3039 (2014).

15. Mou, C. et al. Cryptic Patterning of Avian Skin Confers a Developmental Facility for Loss of Neck Feathering. PLOS Biology 9, e1001028 (2011).

16. Jung, H.-S. et al. Local Inhibitory Action of BMPs and Their Relationships with Activators in Feather Formation: Implications for Periodic Patterning. Dev. Biol. 196, 11–23 (1998).

17. Haupaix, N. et al. The periodic coloration in birds forms through a prepattern of somite origin. Science 361, 6408 (2018).

18. Noramly, S., Freeman, A. & Morgan, B.A. β-catenin signaling can initiate feather bud development. Development 126, 3509–3521 (1999).

19. Turing, A.M. The chemical basis of morphogenesis. Phil. Trans. Roy. Soc. Lond. 237, 37–72 (1952).

20. Gierer, A. & Meinhardt, H. A theory of biological pattern formation. Kybernetik 12, 30–39 (1972).

21. Hillen, T., Painter, K.J. A user’s guide to PDE models for chemotaxis. Journal of Mathematical Biology 58, 183–217 (2009).

22. Corson, F., Couturier, L., Rouault, H., Mazouni, K. & Schweisguth, F. Self-organized Notch dynamics generate stereotyped sensory organ patterns in *Drosophila*. Science 356, eaai7407(2017).

23. Jiang T-X, Jung H-S, Widelitz R, Chuong C-M (1999) Self-organization of periodic patterns by dissociated feather mesenchymal cells and the regulation of size, number and spacing of primordia, Development 126, 4997–5009

24. Shyer, A. E. et al. Emergent cellular self-organization and mechanosensation initiate follicle pattern in the avian skin. Science 357, 811–815 (2017).

25. Hornik, C., Krishan, K., Yusuf, F., Scaal, M. & Brand-Saberi, B. cDermo-1 misexpression induces dense dermis, feathers, and scales. Dev. Biol. 277, p42–50 (2005).

26. Gong, H. et al. Skin transcriptome reveals the dynamic changes in the Wnt pathway during integument morphogenesis of chick embryos. PLOS ONE 13, e0190933 (2018).

27. Henrique, D. et al. Expression of a Delta homologue in prospective neurons in the chick. Nature 375, 787–790 (1995).

28. Hecht, F. New development in FreeFem++. Journal of numerical mathematics 20, 251–266 (2012).

