## Supplementary Information for "Symmetry breaking in the embryonic skin triggers a directional and sequential front of competence during plumage patterning"

### SUPPLEMENTARY FIGURES AND LEGENDS

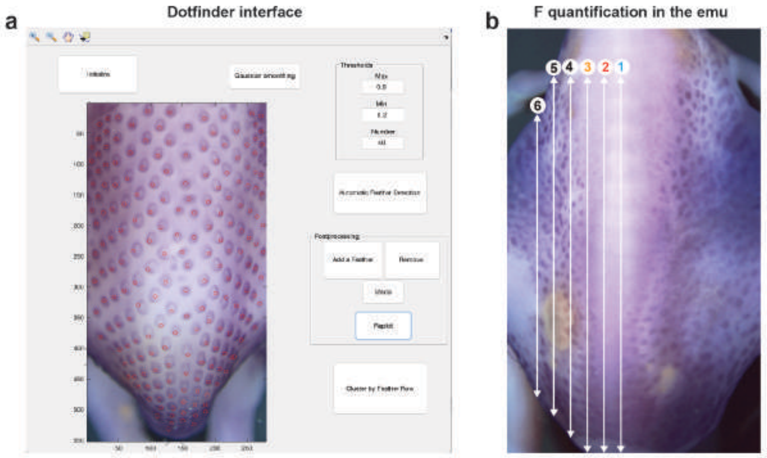


**Supplementary Figure 1: Dotfinder and quantification of F in the emu**

**(a)** User interface for the custom Matlab program for automatic quantification of feather follicles in pictures of whole embryos stained with *ß-catenin* (a Japanese quail at E7,5 is shown), flat skins, and dots in simulation results. The software allows threshold adjustment of the image, automatic detection, post-processing of false positives and negatives, and clustering of follicle/dot rows. **(b)** In the emu, the mean number of feathers per row (F) (i.e., comparable to fr#1-6 in other species) was counted along virtual, equally spaced lines extending from wings to tail (here, at E17).

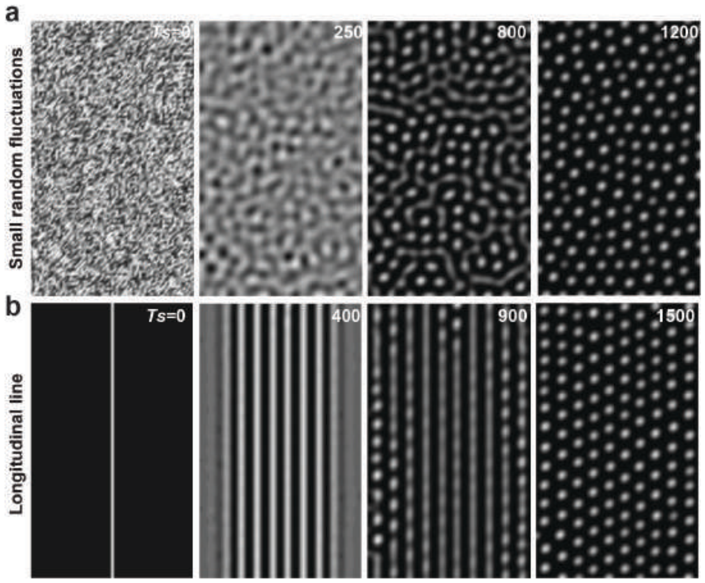

**Supplementary Figure 2: Simulations of a reaction-diffusion model**

**(a)** Simulations of a reaction-diffusion model[13] produce a dotted pattern when initiated on small random fluctuations. **(b)** When this model is forced onto an initial longitudinal line, it produces stripes that once all present, divide into dots to create a dotted motif. **Ts**, simulation time.

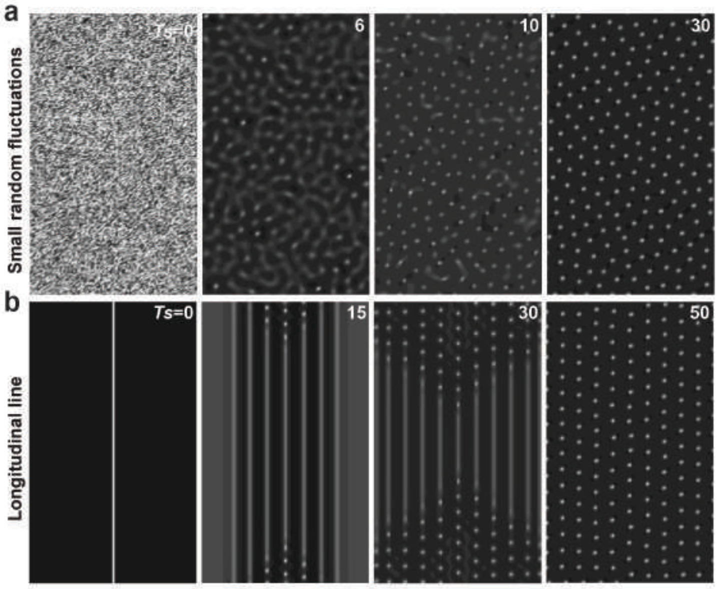


**Supplementary Figure 3: Simulations of a chemotaxis model**

**(a)** Simulations of a chemotaxis model[21] produce dots that appear simultaneously across the whole frame when initiated on small random fluctuations. **(b)** When this model is forced onto an initial longitudinal line, it produces stripes simultaneously that divide into dots, starting in the anterior and posterior regions first and travelling towards the center, resulting in a dotted motif. **Ts**, simulation time.

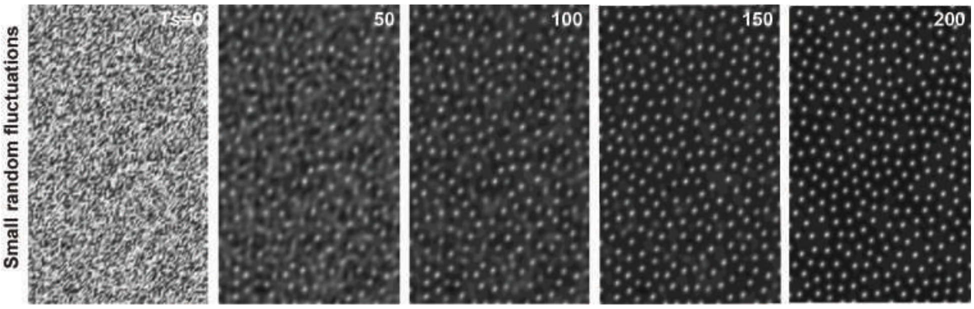


**Supplementary Figure 4: Simulations of the unified model**

Simulations of the unified model (with parameters described in Supp. Table 5) produce dots that appear simultaneously across the whole frame when initiated on small random fluctuations. **Ts**, simulation time.

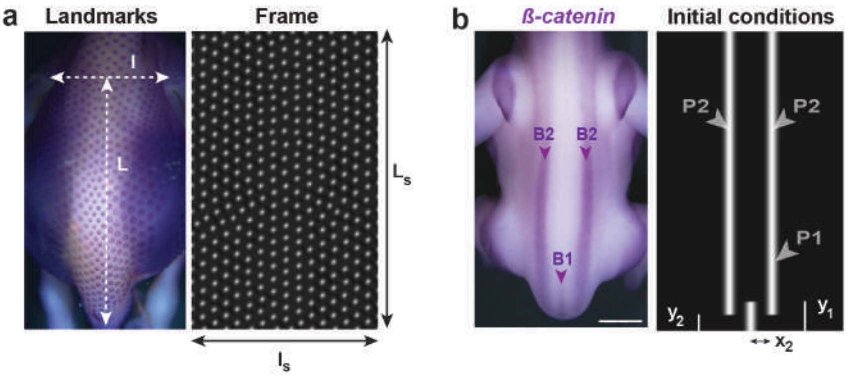


**Supplementary Figure 5: Calculation of frame size and initial conditions**

**(a)** The size of each simulation frame was defined so that its abscissae $l_s$ corresponds to the distance between wings l and its ordinate $L_s$ to the distance between wings and tails L (as shown here on a domestic chicken embryo at E8.5), and reported to obtain F quantified when the model reaches an equilibrium. The obtained ratios for l/ $l_s$ and L/ $L_s$ were then used to define the size of simulation frames for the other four species (all values are shown in Supp. Table 2). **(b)** Initial conditions for P1 and P2 corresponding to measurements of the central location of B1 and B2 *ß-catenin* bands relative to body landmarks and the ratios reported in the Japanese quail are defined by $x_2$ and $y_1$, $y_2$, respectively.

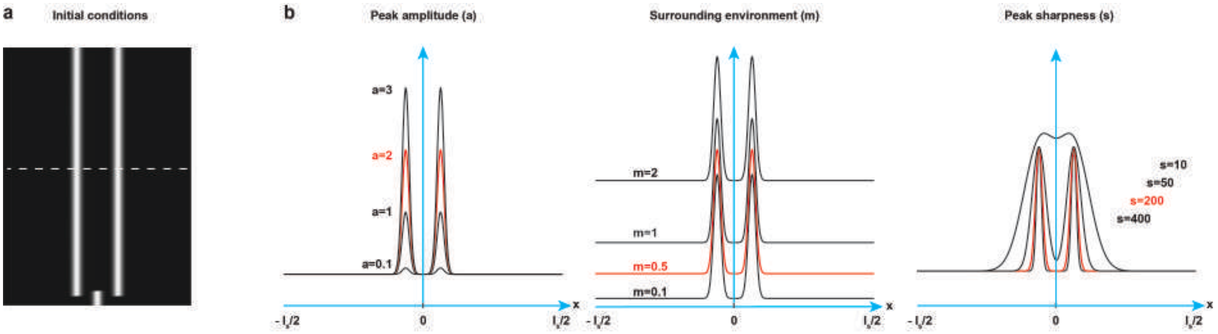


**Supplementary Figure 6: Schematics of Gaussian parameters**

**(a)** Initial conditions for P1 and P2 (see Extended Data Fig. 5). **(b)** Schematics showing the effect of variation in amplitude *a* (left panel), density outside of peaks *m* (middle panel), and sharpness **s** (right panel), compared to reference values (in red), on Gaussian curves corresponding to equation **(2)** at the level of P2 (white dotted line in a). Axes are shown in blue; 0 represents the dorsal midline; $\mathbf{l_s}$: frame width; **x**, abscissae.

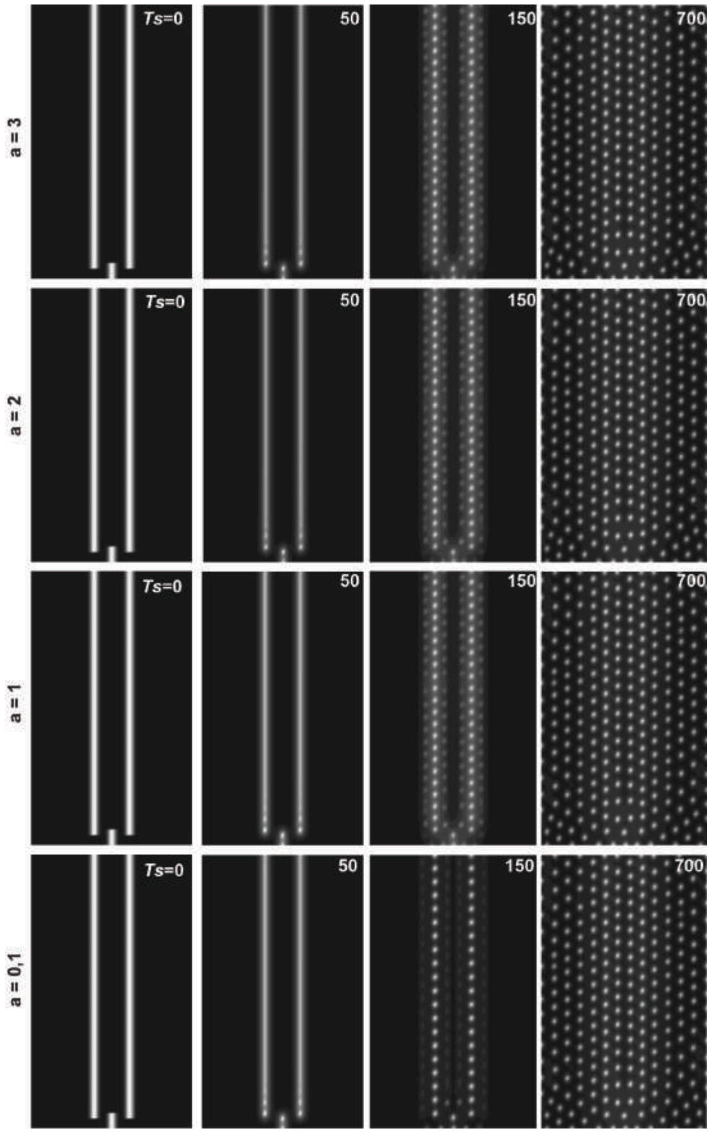


**Supplementary Figure 7: Simulations with variation of the *a* parameter**

Simulations of the unified model with initial conditions corresponding to the Japanese quail, and various amplitude of Gaussian peaks *a* (other parameters equal; the reference parameter is a=2) produce dots in a row-by-row sequence with identical times of simulation (*Ts*=700).

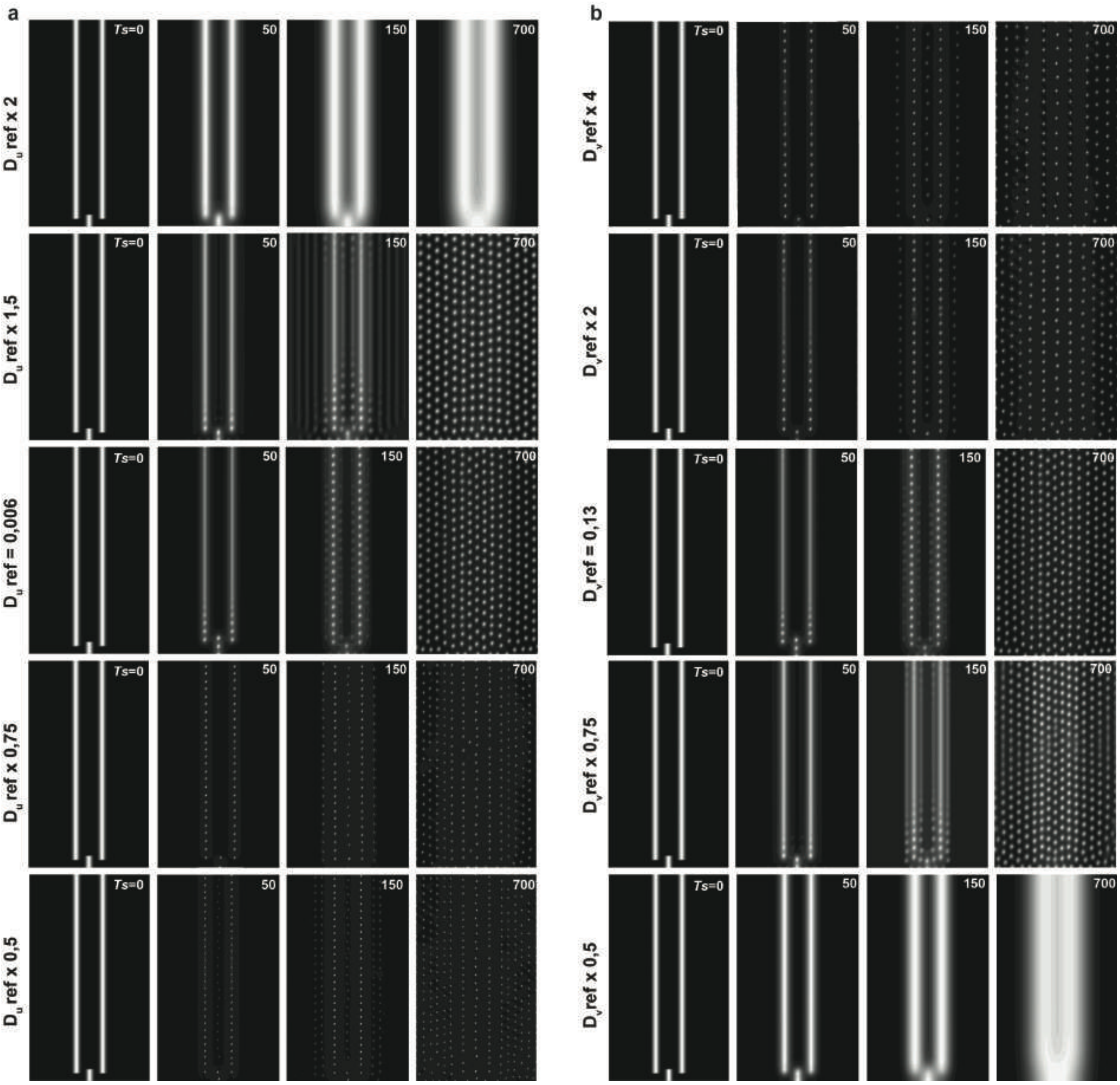


**Supplementary Figure 8: Simulations with variation of $D_u, D_v$ parameters**

Simulations of the unified model with initial conditions corresponding to the Japanese quail, and various diffusivity of the activator $D_u$ **(a)** or of the inhibitor $D_v$ **(b)**, other parameters equal (references parameters are $D_u$ ref = 0,006 and $D_v$ ref = 0,13) produce dots varying in size and spacing but in a row-by-row sequence. **Ts**, simulation time.

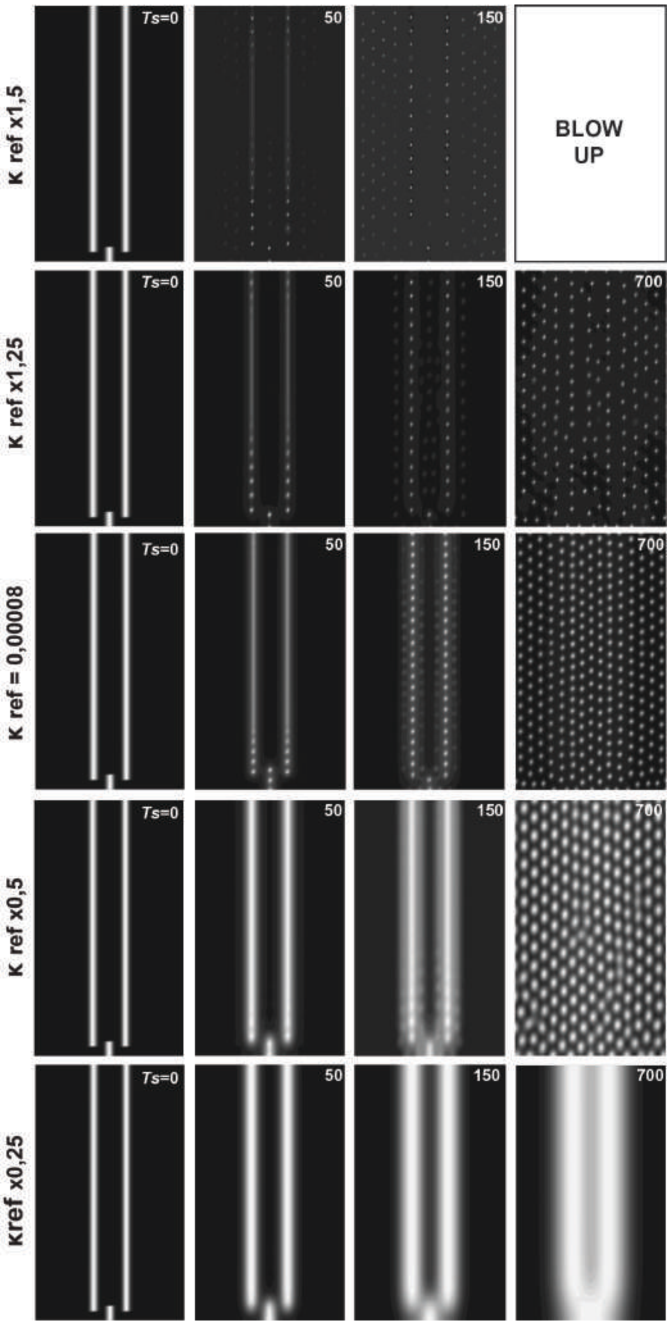


**Supplementary Figure 9: Simulations with variation of the $\kappa$ parameter**

Simulations of the unified model with initial conditions corresponding to the Japanese quail, and various diffusivity of the sensitivity to chemotaxis $\kappa$ (other parameters equal, references parameters are $\kappa$ ref = 0,00008) produce dots varying in size and spacing but in a row-by-row sequence. **Ts**, simulation time.

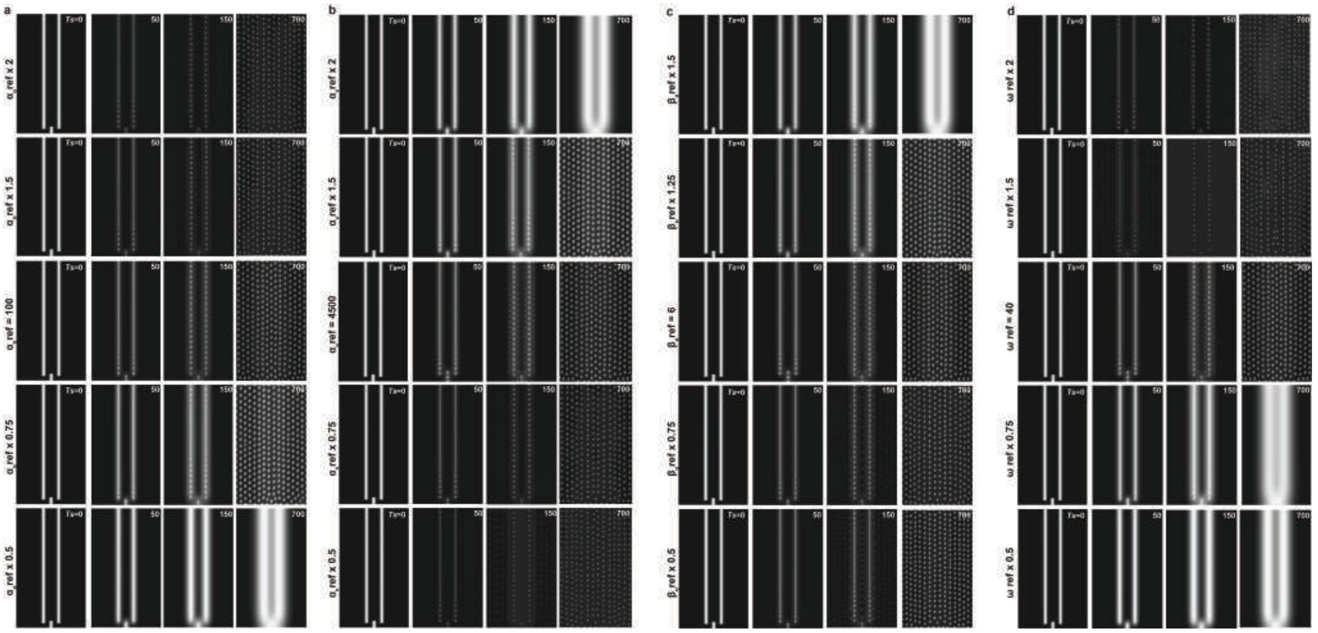

**Supplementary Figure 10: Simulations with variation of the $\alpha_u, \alpha_v, \beta_u$ and $\omega$ parameters**

Simulations of the unified model with initial conditions corresponding to the Japanese quail, and various production rates of the activator (or repressor) by the cells $\alpha_u$ ($\alpha_v$), the saturation threshold $\beta_u$ and autocatalysis sensitivity $\omega$ of the activator (other parameters equal, references parameters are $\alpha_u$ ref = 100, $\alpha_v$ ref = 4500, $\beta_u$ ref = 6, and $\omega$ ref = 40) produce dots varying in size and spacing but in a row-by-row sequence. **Ts**, simulation time.

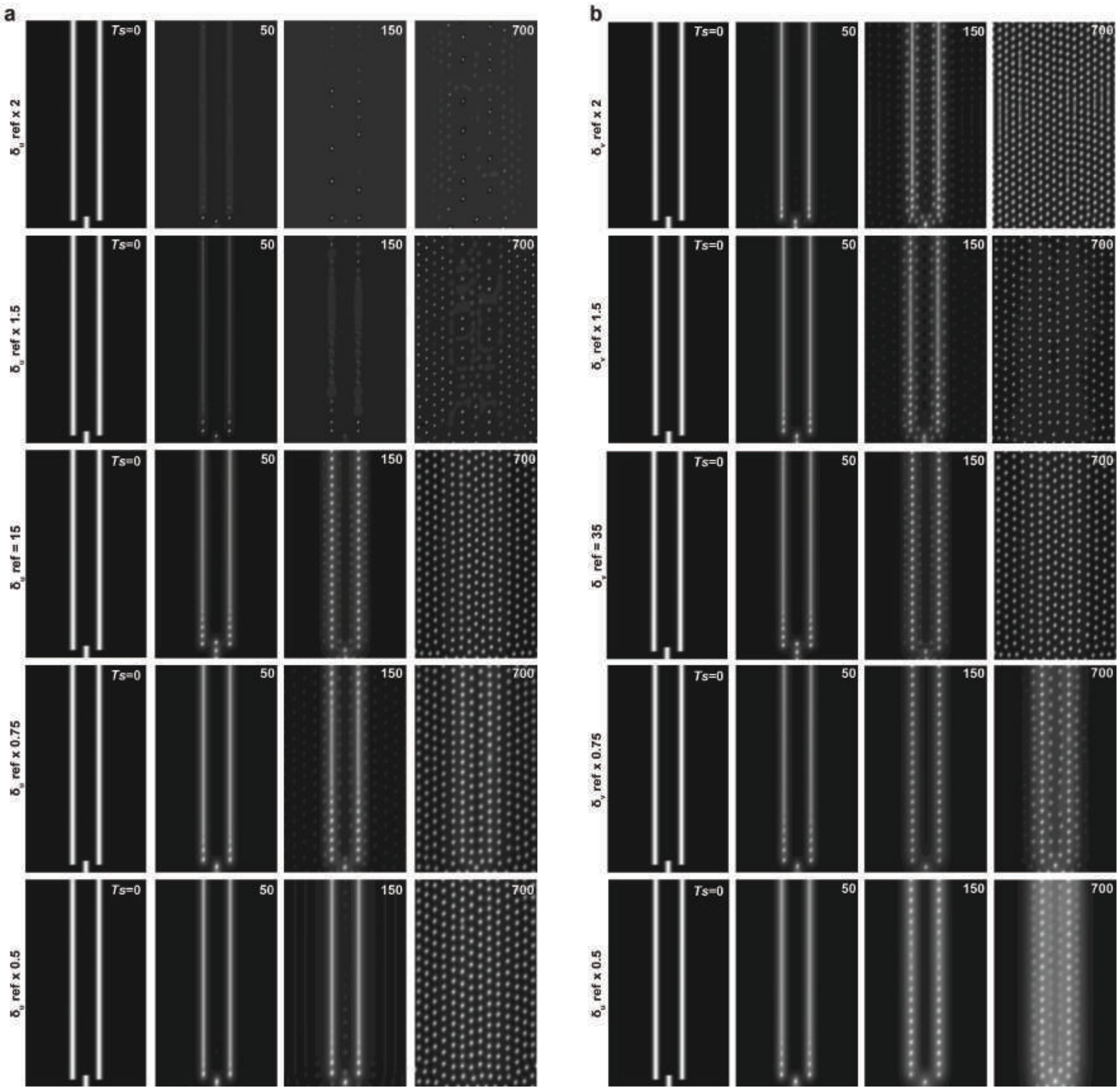


**Supplementary Figure 11: Simulations with variation of the $\delta_u$, $\delta_v$ parameters**

Simulations of the unified model with initial conditions corresponding to the Japanese quail, and various degradation rates of the activator (or repressor) $\delta_u$ ($\delta_v$), other parameters equal (references parameters are $\delta_u$ ref = 15, $\delta_v$ ref = 35) produce dots varying in size and spacing but in a row-by-row sequence. **Ts**, simulation time.

### SUPPLEMENTARY TABLES

#### Supplementary Table 1: Number of embryos assessed for *ß-catenin* expression

| **Stage (E)** | Stage 1 | Stage 2 | Stage 3 | Stage 4 | Stage 5 |
|---|---|---|---|---|---|
| *Gallus gallus* | 2 | 3 | 3 | 4 | 6 |
| *Coturnix japonica* | 5 | 4 | 2 | 4 | 4 |
| *Phasianus colchicus* | 5 | 3 | 3 | 4 | 5 |
| *Taeniopygia guttata* | 4 | 4 | 6 | 7 | 9 |
| *Dromaius novaehollandiae* | 2 | 1 | 1 | 1 | 1 |

#### Supplementary Table 2: *In vivo* measurements and *in silico* domain units

| | *G. gallus* | *C. japonica* | *P. colchicus* | *T. guttata* | *D. novaehollandiae* |
|---|---|---|---|---|---|
| Wings-tail distance L (mm) | 11 | 10 | 10.1 | 6.8 | 13.5 |
| Wing width l (mm) | 5 | 4.7 | 4.8 | 2.5 | 7 |
| *In silico* domain size $L_s$ x $l_s$ | 5 x 3.5 | 4.5 x 3.3 | 4.6 x 3.4 | 3.1 x 1.8 | 6.1 x 4.9 |

#### Supplementary Table 3: Parameters of initial conditions of simulation

| **Parameters** | *G. gallus* | *C. japonica* | *P. colchicus* | *T. guttata* |
|---|---|---|---|---|
| m | 0.5 | | | |
| a | 2 | | | |
| $s_m$ | 200 | 200 | 20 | 400 |
| $s_l$ | 200 | 200 | 200 | 400 |
| $x_l$ | 0.25 | 0.36 | 0.25 | 0.2 |
| $y_l$ | 2.6 | 0.1 | 0.1 | 0.9 |
| $y_m$ | 2.8 | 0.3 | 2 | 0.85 |

**Supplementary Table 4: Tested reaction-diffusion models**

| |
|---|
| $\begin{cases}\partial_t \mathrm{u} = D_u \Delta u + \min(\max(a_u u + b_u v + c_u, 0), F_{max}) \\ \partial_t \mathrm{v} = D_v \Delta v + \min(\max(a_v u + b_v v + c_v, 0), G_{max})\end{cases}$ |
| $\begin{cases}\partial_t \mathrm{u} = D_u \Delta u + p_u \dfrac{u^2}{(1 + k_2 v)({k_1}^2 + u^2)} + \alpha_u - \delta_u u \\ \partial_t \mathrm{v} = D_v \Delta v + p_v \dfrac{u^2}{{k_3}^2 + u^2} + \alpha_v - \delta_v v\end{cases}$ |
| $\begin{cases}\partial_t \mathrm{u} = D_u \Delta u + p_u \dfrac{u^2}{(\gamma + v)(1 + k u^2)} - \delta_u u \\ \partial_t \mathrm{v} = D_v \Delta v + p_v \dfrac{u^2}{(\gamma + v)(1 + k u^2)} - \delta_v v\end{cases}.$ |
| $\begin{cases}\partial_t \mathrm{u} = D_u \Delta u - u v^2 + \alpha_u - \delta_u u \\ \partial_t \mathrm{v} = D_v \Delta v + u v^2 - \delta_v v\end{cases}.$ |
| $\begin{cases}\partial_t \mathrm{u} = D_u \Delta u - \dfrac{4uv}{1 + v^2} + \alpha_u - u \\ \partial_t \mathrm{v} = D_v \Delta v + \alpha_v (u - \dfrac{uv}{1 + v^2})\end{cases}.$ |

**Supplementary Table 5: Reference parameters of the unified model**

| Parameter | **Value** | **Biological interpretation** |
|---|---|---|
| $D_n$ | $7*10^{-5}$ | Cell diffusion rate |
| $D_u$ | $6*10^{-3}$ | Activator's diffusion rate |
| $D_v$ | 0.13 | Inhibitor diffusion rate |
| $\kappa$ | $8*10^{-5}$ | Chemotaxis sensitivity |
| $\delta_u$ | 15 | Activator degradation rate |
| $\delta_v$ | 35 | Inhibitor degradation rate |
| $\alpha_u$ | 100 | Activator production rate by cells |
| $\omega$ | 40 | Activator autocatalysis sensitivity |
| $\beta_u$ | 6 | Activator saturation threshold |
| $\alpha_v$ | 4500 | Inhibitor production rate by cells |
| $\alpha$ | $1*10^{-3}$ | Proliferation rate |
| $\beta$ | 3 | Cell density threshold |

**Supplementary Table 6: Tested doses of Colchicine**

| Colchicine concentration (mg/L) | 0.001 | 0.01 | 0.025 | 0.05 | 0.1 | 0.2 | 0.5 | 1 | 2 | 4 | 40 |
|---|---|---|---|---|---|---|---|---|---|---|---|
| Lethality | no | no | no | no | yes | yes | yes | yes | yes | yes | yes |
